# Clinical, socio-demographic, and parental correlates of early autism traits in a community cohort

**DOI:** 10.1101/2022.09.26.508121

**Authors:** Oliver Gale-Grant, Andrew Chew, Shona Falconer, Lucas G.S França, Sunniva Fenn-Moltu, Laila Hadaya, Nicholas Harper, Judit Ciarrusta, Tony Charman, Declan Murphy, Tomoki Arichi, Grainne McAlonan, Chiara Nosarti, A David Edwards, Dafnis Batalle

## Abstract

**Background:** Autism traits emerge between the ages of 1 and 2. It is not known if experiences which increase the likelihood of childhood autism are related to early trait emergence, or if other exposures are more important. Identifying factors linked to toddler autism traits in the general population may improve our understanding of the mechanisms underlying atypical neurodevelopment.

**Methods:** Clinical, socio-demographic, and parental information was collected at birth from 536 toddlers in London, UK (gestational age at birth, sex, maternal body mass index, age, parental education level, parental first language, parental history of neurodevelopmental disorders) and at 18 months (parent cohabiting status, two measures of social deprivation, three measures of maternal parenting style, and a measure of maternal postnatal depression). General neurodevelopment was assessed with the Bayley Scales of Infant and Toddler Development, 3^rd^ Edition (BSID-III), and autism traits were assessed using the Quantitative Checklist for Autism in Toddlers (Q-CHAT). Multivariable models were used to identify associations between variables and Q-CHAT. A model including BSID-III was used to identify factors associated with Q-CHAT independent of general neurodevelopment. Models were also evaluated addressing variable collinearity with principal component analysis (PCA).

**Results:** A multivariable model explained 20% of Q-CHAT variance, with four individually significant variables (two measures of parenting style and two measures of socio-economic deprivation). After adding general neurodevelopment into the model 36% of Q-CHAT variance was explained, with three individually significant variables (two measures of parenting style and one measure of language development). After addressing variable collinearity with PCA, parenting style and social deprivation were positively correlated with Q-CHAT score via a single principal component, independently of general neurodevelopment. Neither sex nor family history of autism were associated with Q-CHAT score.

**Limitations:** The Q-CHAT is parent rated and is therefore a subjective opinion rather than a clinical assessment. We measured Q-CHAT at a single timepoint, and to date no participant has been followed up in later childhood, so we are focused purely on emerging traits rather than clinical autism diagnoses.

**Conclusions:** Autism traits are common at age 18 months, and greater emergence is specifically related to exposure to early life adversity.

## Background

Autism spectrum disorders (ASD) are typically diagnosed between 4 and 7 years of age (1, 2). The age at symptom onset however is often lower than this, with neurodivergence first being suspected by parents in most instances between 1 and 2 years of age (3). Screening tools aiming to quantify autism traits in this age group are well established and cut-off points with high sensitivity (albeit at the cost of low specificity (4)) for predicting a future clinical autism diagnosis have been demonstrated (5, 6), although results in some real-world cohorts are less promising (7). One such tool is the Qualitative Checklist for Autism in Toddlers (Q-CHAT) (8). The Q-CHAT is a 25-item questionnaire with each item rated by the parents from 0-4. It has been validated for use in multiple countries (9-13), and has a positive predictive value of 28% for a future ASD diagnosis (using screening at two timepoints) (14). Autistic traits exist in the population as a continuum (15), and most individuals screened, typically developing or otherwise, will display at least some autism traits at age 18 months (16).

The likelihood of receiving an autism diagnosis is associated with both genetic and environmental factors (17, 18), and the same may be true of early autism traits. Some factors are known to correlate with autism traits at age 18 months – for example, sex (with males scoring higher than females) (7, 8, 19, 20) or preterm birth (10, 21). However, beyond these factors there is a relative lack of research into what else may influence the emergence of autism traits in early life, although single studies have linked maternal nausea and vomiting during pregnancy (22), neonatal illness (23), maternal depression and anxiety (24, 25), immigrant mothers (26) and lower levels of parental education (24) with higher scores on 18 month autism screening tools. Q-CHAT score at 18 months has also been shown to be negatively correlated with general language development (10). The broader developmental phenotype is known to be influenced by a wide range of exposures, including preterm birth (27), neonatal illness (28), and multiple psychosocial factors (29-32). Given that Q-CHAT is known to correlate with general language development, it is reasonable to hypothesise that Q-CHAT scores may themselves be influenced by these same exposures.

As well as research using structured tools there are previous studies which examine exposures associated with single features of social communication development in toddlerhood. Multiple factors including less responsive or less effective maternal parenting styles (33, 34), greater maternal depression and experience of trauma (35) and a lower quality home environment (36) have been correlated with less favourable social communication development in toddlerhood.

Because greater autism trait emergence at age 18 months is associated with a greater likelihood of childhood autism (14) understanding correlates of the Q-CHAT score at 18 months may help us to understand what early life experiences are associated with an increased likelihood of a future autism diagnosis in some individuals. The developing Human Connectome Project (dHCP) has collected Q-CHAT scores, other neurodevelopmental measures and demographic information from a large cohort of 18-month-old toddlers in London, UK. Using this dataset, we aimed to characterise correlates of Q-CHAT score. We hypothesised that, in keeping with the known associations between early life adversity and other measures of neurodevelopment, we would observe a pattern of psychosocial adversity being associated with higher Q-CHAT scores. Relationships between variables and Q-CHAT score are presented in both univariable (in part to inform future studies which may only have some of our variables available) and multivariable models, with scores from the Bayley Scales of Infant and Toddler Development, 3^rd^ Edition (BSID-III) (37) additionally included as a covariate to understand whether any relationships between these early life experiences and autism traits are influenced by general neurodevelopment.

## Methods

### Sample

This study is based on a sample of neonates participating in the Developing Human Connectome Project (dHCP, http://www.developingconnectome.org/). Participants were all recruited at St Thomas’ Hospital, London, UK. There were no specific inclusion or exclusion criteria for enrolment in this study, and recruitment was primarily from the antenatal clinic with no specific stratification.

Toddlers were invited for neurodevelopmental assessment at 18 months post-expected delivery date; appointments were made according to family availability as close as possible to this time-point. The only inclusion criteria for this manuscript from the overall cohort was completion of the neurodevelopmental assessment. There were no exclusion criteria.

This project has received UK NHS research ethics committee approval (14/LO/1169, IRAS 138070), and conducted in accordance with the World Medical Association’s Code of Ethics (Declaration of Helsinki). Written informed consent was obtained from parents at recruitment into the study.

### Data Collection

Data collection took place either at St Thomas’ Hospital, London, UK, or via questionnaires distributed to the participants’ parents. At the time of birth, clinical variables, gestational age at birth and sex were extracted from the medical records of participants in the study; and maternal age, maternal pre-pregnancy BMI, and parent ASD/attention deficit hyperactivity disorder (ADHD) history were also collected via a maternal questionnaire. The last of these was asked in the format “Have you or your child’s biological father ever been diagnosed with Attention Deficit Hyperactivity Disorder (ADHD) or Autism?” This was a yes/no question.

At the time of birth, the socio-demographic status of participant families was recorded as measured by the Index of Multiple Deprivation Rank (IMD), a postcode-based score assigned to every address in the UK which gives a composite measure of socio-economic disadvantage, based on the mother’s address at the time of birth. A lower score corresponds to greater geographical deprivation, with 1 being the lowest score possible and 32,844 being the highest score possible.

Further socio-demographic information was collected by questionnaire: maternal age at leaving education (“At what age were you last in full time education?”), maternal 1^st^ language (“Is English your first language”?), and parent cohabiting status. The Cognitively Stimulating Parenting Scale (CSPS), a questionnaire assessing the availability of resources to support cognitively stimulation parenting, associated to both parenting style and socio-economic deprivation was also collected (38, 39). The CSPS was updated to include items relating to access to mobile phones and apps. A higher score is indicative of a more stimulating home environment, with a minimum possible score of 0 and a maximum possible score of 40. The Q-CHAT score (a parent reported questionnaire) was collected at the time of 18-month follow-up. This gives a score between 0 and 100, with higher scores indicative of more autism traits. The Bayley Scales of Infant and Toddler Development, 3^rd^ edition (BSID-III) (37), was administered by either a Chartered Psychologist or Paediatrician when the children were 18-months of age. The composite scores (Cognitive, Motor and Language) are used for analysis in this study. Two measures of parenting style were also collected at this time. The first of these, the Parenting Scale (40), is a self-reported tool that measures three different dimensions of parenting: Laxness, the tendency to behave passively and give in to misbehaviour; Over-reactivity, which measures anger, meanness and irritability in parenting; and Verbosity, a measure of parental dependence on talking even when ineffective as a discipline style. The dimensions have a minimum score of 1, and a maximum of 7. The Edinburgh Postnatal Depression Scale (EPDS) was also completed at follow-up. This is a well-established self-reported tool for quantifying postnatal depressive symptoms, with a minimum score of 0 and a maximum of 30. Higher scores are indicative of more depressive symptoms (41).

### Statistical Analysis

Univariable associations between variables and Q-CHAT score were tested by Pearson’s correlation or t-test as appropriate. Associations between variables of interest and Q-CHAT score were assessed by generalized linear model (GLM). Statistical significance was tested with random permutation tests, using 10,000 permutations. P-values are reported uncorrected, with those surviving multiple comparisons via false discovery rate (FDR) indicated (42). Principal component analysis was used to characterize the latent structure of independent variables, and to address collinearity between linear variables. The “elbow method” (43) was used to determine the optimal number of principal components (PCs) to use in later analyses. Associations between PC scores and the original input variables was determined by Pearson’s correlation, with p<0.05 after FDR correction considered significant.

Analyses were performed and figures made using Rstudio v4.0.2 (Rstudio, MA, U.S.A). The “FDRestimation”, and “corrplot” packages were additionally used (44, 45). PCA was performed using the “prcomp” function from base R rather than a dedicated package. Our code to implement random permutation tests for GLMs in R can be downloaded from https://github.com/CoDe-Neuro/ptestR.

### Data availability

The dHCP is an open-access project. Data from the project can be downloaded by registering at https://data.developingconnectome.org. Analyses presented here include data to be included in future releases.

## Results

### Population

At the time of the study commencing, 644 individuals in the dHCP dataset had a Q-CHAT score available. Of these 536 had a complete set of demographic data and were included in the study. A comparison between individuals included and excluded is shown in Supplementary Table S1. There were some differences between those included and excluded – individuals included in the study experienced on average lower geographical deprivation (IMD Rank), lower maternal depression, and less extreme parenting styles. The characteristics of the sample used, and the univariate relationships of each variable to Q-CHAT score are shown in Table 1. A frequency distribution of Q-CHAT scores is in Supplementary Figure S1.

**Table 1.** Sample Characteristics. Mean, standard deviation, and range displayed for linear variables. Frequency displayed for categorical variables. Correlations to QCHAT calculated by Pearson’s r or t-test as appropriate. Significant univariable correlations are shown in bold, those remaining significant after FDR correction indicated by *.

| **Outcome variable** |  |  |
| --- | --- | --- |
| Total Q-CHAT Score, *Mean (SD), Range* | 30.1 (5.9), 8-70 | - |
| **Clinical variables** |  | ***r* (p)** |
| Age at follow-up [months], *Mean (SD), Range* | 18.8 (1.6), 16-26 | 0.010 (0.535) |
| Gestational Age at Birth [weeks], *Mean (SD), Range* | 38.1 (3.9), 23.0-43.0 | -0.067 (0.120) |
| BMI [kg/m<sup>2</sup>], *Mean (SD), Range* | 24.2 (4.4), 15.3-43.4 | **0.093 (0.030)** |
| Mother Age [years], *Mean (SD), Range* | 34.3 (4.7), 17-52 | **-0.105 (0.014)** |
|  |  | ***t* (p)** |
| Sex [Male (0), Female (1)], *N (%)* | 278 (52%), 258 (48%) | 1.820 (0.068) |
| Parent ASD/ADHD Diagnosis [Yes (1), No (0)], *N (%)* | 28 (5%), 508 (95%) | -0.4246 (0.674) |
| **Socio-demographic variables** |  | ***r* (p)** |
| IMD Rank, *Mean (SD), Range* | 14626.2 (7409.2), 2410-32726 | **-0.190 (<0.001)\*** |
| CSPS, *Mean (SD), Range* | 20.5 (3.5), 7-28 | **-0.219 (<0.001)\*** |
| Mother Education [years], *Mean (SD), Range* | 23.6 (4.5), 12-43 | 0.001 (0.957) |
|  |  | ***t* (p)** |
| Mother 1<sup>st</sup> Language [English (1), Other (0)], *N (%)* | 338 (63%), 198 (37%) | **4.518 (<0.001)\*** |
| Parents Cohabiting [Yes (0), No (1)], *N (%)* | 520 (97%), 16 (3%) | -1.650 (0.119) |
| **Parental-psychological variables** |  | ***r* (p)** |
| Mother Laxness, *Mean (SD), Range* | 2.9 (0.8), 1-5.6 | **0.286 (<0.001)\*** |
| Mother Over-reactivity, *Mean (SD), Range* | 2.2 (0.7), 1-5.1 | **0.180 (<0.001)\*** |
| Mother Verbosity, *Mean (SD), Range* | 3.4 (0.8), 1-6.4 | **0.300 (<0.001)\*** |
| Mother EPDS, *Mean (SD), Range* | 4.5 (4.2), 0-28 | **0.127 (<0.001)\*** |
| BSID-III Cognitive Composite, *Mean (SD), Range* | 101.0 (11.1), 55-130 | **-0.358 (<0.001)\*** |
| BSID-III Language Composite, *Mean (SD), Range* | 98.2 (15.4), 47-153 | **-0.528 (<0.001)\*** |
| BSID-III Motor Composite, *Mean (SD), Range* | 101.5 (10.2), 52-130 | **-0.267 (<0.001)\*** |

Nine variables were significantly associated with Q-CHAT score. BMI (r=0.093, p=0.030), EPDS (r=0.127, p<0.001), and three measures of maternal parenting style; laxness (r=0.286, p<0.001), over-reactivity (r=0.180, p<0.001) and verbosity (r=0.300, p<0.001) were positively correlated with Q-CHAT score, and mother’s age (r=-0.105, p=0.014), IMD rank (r=-0.190, p<0.001) and CSPS score (r=-0.219, p<0.001) were negatively correlated with Q-CHAT score. Total Q-CHAT scores were significantly higher in individuals whose mother’s spoke a language other than English as their 1^st^ language (t=4.52, p<0.001).

All BSID-III composite scores were negatively associated with Q-CHAT score. The strongest association was with Language Composite Score (r=-0.528, p<0.001).

### Multivariable models of Q-CHAT score

We assessed the association of all variables with Q-CHAT score in two separate multivariable models, with or without the addition of BSID-III composite scores to control for the effect of general neurodevelopment and identify specific relationships between demographic variables and Q-CHAT score.

**Table 2.** General linear model of the association between clinical, socio-demographic, and parental variables and Q-CHAT with or without the addition of BSID-III Cognitive, Motor and Language Composite Scores to the model. Non-reference categories are as follows: Sex – Male, Parent ASD/ADHD Diagnosis – Yes, Mother 1^st^ Language – Not English.

|  | *Without BSID-III*<br>*r<sup>2</sup> (Adj. r<sup>2</sup>) = 0.20(0.19), p<0.001* |  | *With BSID-III*<br>*r<sup>2</sup> (Adj. r<sup>2</sup>) = 0.36(0.34), p<0.001* |  |
| --- | --- | --- | --- | --- |
|  | **t** | **p** | **t** | **p** |
| Gestational Age at Birth | -1.94 | 0.052 | -0.45 | 0.646 |
| BMI | 0.57 | 0.567 | 0.95 | 0.341 |
| Mother Age | -0.73 | 0.461 | -1.17 | 0.239 |
| Sex | **-2.32** | **0.020** | -0.83 | 0.407 |
| Parent ASD/ADHD Diagnosis | 0.57 | 0.567 | -0.23 | 0.813 |
| IMD Rank | **-2.56** | **0.010\*** | **-2.06** | **0.039** |
| CSPS | **-3.38** | **<0.001\*** | -1.17 | 0.238 |
| Mother Education | -0.08 | 0.929 | 0.35 | 0.719 |
| Mother 1<sup>st</sup> Language English | -1.58 | 0.113 | -0.01 | 0.994 |
| Parents Cohabiting | 1.74 | 0.082 | 1.49 | 0.135 |
| Mother Laxness | **3.79** | **<0.001\*** | **2.68** | **0.007\*** |
| Mother Over-reactivity | 1.11 | 0.269 | 1.04 | 0.297 |
| Mother Verbosity | **3.29** | **0.001\*** | **3.39** | **<0.001\*** |
| Mother EPDS | 1.72 | 0.08 | 1.26 | 0.206 |
| Cognitive Composite | NA | NA | -0.28 | 0.779 |
| Language Composite | NA | NA | **-8.32** | **<0.001\*** |
| Motor Composite | NA | NA | -0.41 | 0.677 |

*Bold indicates p<0.05, \* indicates significance after FDR multiple comparison correction (α<0.05)*

A multivariable model without BSID-III explained 20% of Q-CHAT variance. After FDR correction four variables were individually associated with Q-CHAT score: IMD Rank (t=-2.56,p=0.010) and CSPS (t=-3.38,p<0.001) were negatively associated and Mother Laxness (t=3.79,p<0.001) and Mother Verbosity (t=3.29, p=0.001) were positively associated. After adding BSID-III composite scores to the model two of these (Mother Laxness and Mother Verbosity) remained significantly associated with Total Q-CHAT score (t=2.68,p=0.007 and t=3.39,p<0.001 respectively), in addition to BSID-III language composite score (t=-8.32, p<0.001), which was negatively associated with Total Q-CHAT score. Notably sex and parent ASD/ADHD diagnosis status did not correlate individually with Q-CHAT score in either model.

**Figure 1.**
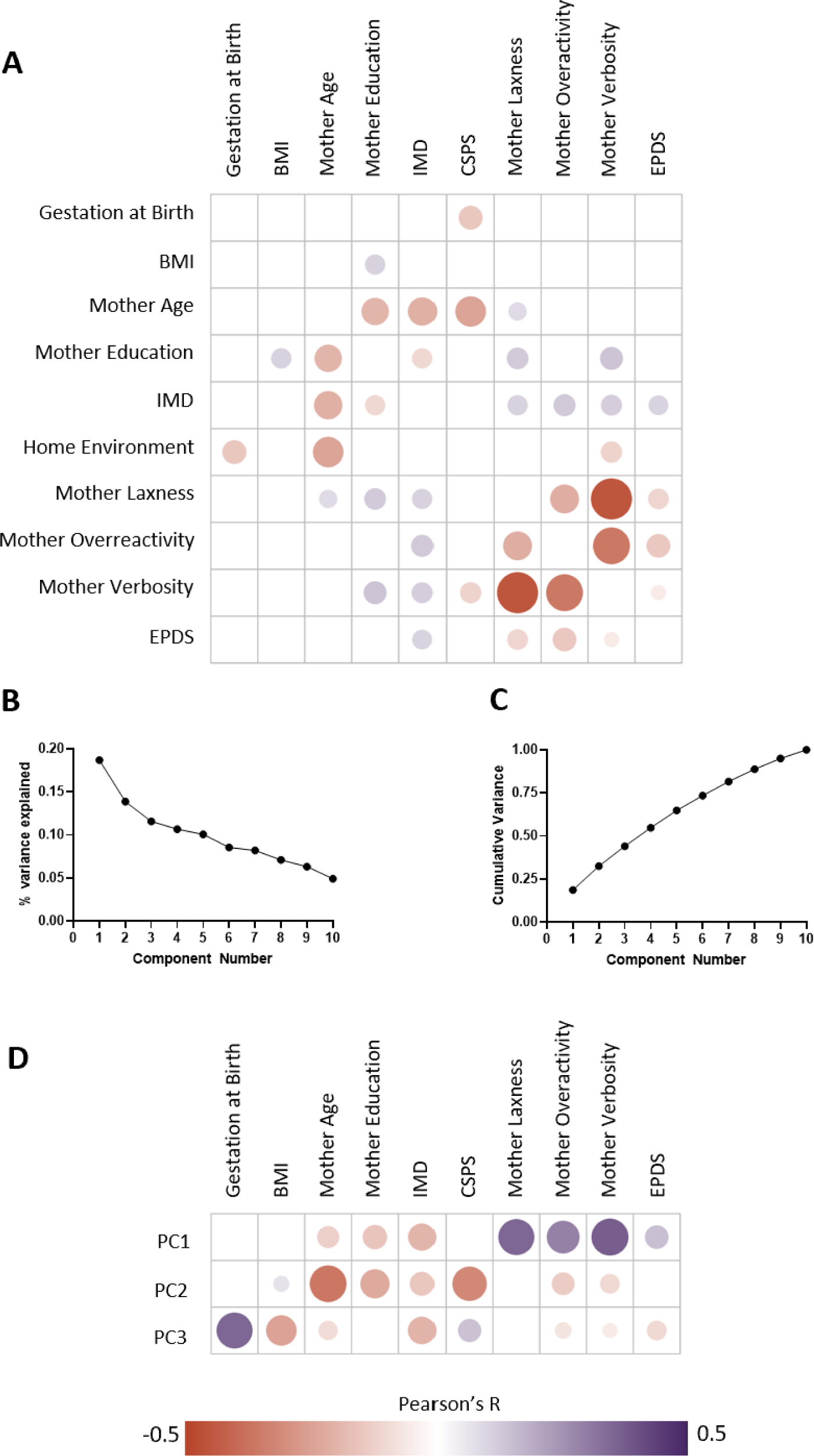
Principal Component Analysis of linear variables. **A** Correlogram of associations between linear variables. Pearson’s r indicated for correlations with p<0.05. **B** Scree plot of PCA components **C** Cumulative variance plot of PCA components **D** Correlations of original linear variables to principal components. Correlation indicated by size and colour of circle. Only correlations remaining significant (p<0.05) after FDR correction are shown. Values of each correlation are shown in Supplementary Table S2.

A limitation of interpreting these models is the collinearity between demographic variables (Figure 1A). In order to address this without removing variables from the model, we performed a PCA of the linear variables to obtain orthogonal components, which we then used in a general linear model in place of the original linear variables (46). We selected the first 3 principal components (PCs) to represent our data (Figure 1B). The multivariable models associating demographic variables and BSID-III composite scores with Q-CHAT score (Table 2) were subsequently repeated, with linear variables being replaced by PCA components 1-3 (Table 3). Details of variable correlations with each PC are shown in Figure 1D. PC1 captures variable associations which are associated with positive parenting styles and low socio-economic deprivation, PC2 is associated with socio-economic deprivation and a less stimulating home environment, and PC3 is associated with low clinical adversity.

17% of Q-CHAT variance was explained by a model including 3 PCs and the categorical variables only, with only PC1 remaining statistically significant in the model after FDR correction (t=-8.17, p<0.001, Table 3). 36% of Q-CHAT variance was explained by the model including BSID-III scores to account for general neurodevelopment, with PC1 and BSID-III language composite scores statistically significant (t=-6.59 and t=-8.96 respectively, p<0.001, Table 3).

**Table 3.** General linear model of the association between demographic variables, BSID-III composite scores and Q-CHAT. Linear variables were first transformed into orthogonal components via PCA. PC1 captures variable associations which are associated with positive parenting styles and low socio-economic deprivation, PC2 is associated with low socio-economic deprivation and expressive parenting styles, and PC3 is associated with variables describing clinical adversity.

|  | *Without BSID-III* |  | *With BSID-III* |  |
| --- | --- | --- | --- | --- |
|  | *r<sup>2</sup> (Adj. r<sup>2</sup>) = 0.17(0.16), p<0.001* |  | *r<sup>2</sup> (Adj. r<sup>2</sup>) = 0.35(0.33), p<0.001* |  |
|  | **t** | **p** | **t** | **p** |
| PC1 | **-8.17** | **<0.001\*** | **-6.59** | **<0.001\*** |
| PC2 | -1.86 | 0.062 | -0.70 | 0.480 |
| PC3 | 1.29 | 0.194 | 0.10 | 0.913 |
| Sex | -2.02 | 0.043 | -0.48 | 0.630 |
| Parent ASD/ADHD Diagnosis | 0.59 | 0.550 | -0.26 | 0.790 |
| Mother 1<sup>st</sup> Language English | -1.86 | 0.063 | -0.06 | 0.948 |
| Parents Cohabiting | 2.04 | 0.041 | 1.51 | 0.129 |
| Cognitive Composite | NA | NA | -0.55 | 0.577 |
| Language Composite | NA | NA | **-8.96** | **<0.001\*** |
| Motor Composite | NA | NA | -0.15 | 0.880 |

*Bold indicates p<0.05, \* indicates significance after FDR multiple comparison correction (α<0.05)*

PC1 (positive parenting styles and low socio-economic deprivation) is negatively correlated with Q-CHAT score (t=-8.17, p<0.001) - ie, individuals with more adversity have lower Q-CHAT scores. Via this PC we can see that maternal age at last full-time education, three measures of parenting style and EPDS are positively associated with Q-CHAT score; whereas maternal age at leaving education and IMD rank are negatively associated with Q-CHAT score. Once again it is worth noting that sex and parent ASD/ADHD diagnosis status did not correlate with Q-CHAT score in either model.

## Discussion

We observed correlations of Q-CHAT score with measures of parenting style and measures of socio-demographic adversity, with the former category demonstrating the strongest associations. Conversely, some variables known to increase the likelihood of an autism diagnosis in later childhood, such as male sex (47), a family history of autism (48) and gestational age at birth (49) were not associated with Q-CHAT scores.

A multivariable model of demographic variables explained 20% of Q-CHAT variance. In this model four variables (two measures of social deprivation and two measures of parenting style) were individually significantly associated with Q-CHAT score. Adding measures of general neurodevelopment to this model increased the explained variance to 36%, however this also resulted in two variables, IMD Rank and CSPS (measures of social deprivation) no longer being individually significantly associated with Q-CHAT score. Taken together this suggests that maternal parenting style is specifically associated with Q-CHAT score, whereas that the association of social deprivation with Q-CHAT is partially explained by general neurodevelopment.

Maternal verbosity had the strongest association with Q-CHAT score of any variable tested, remaining significantly associated with Q-CHAT score in multivariable models with and without general neurodevelopment. The mechanism via which this association occurs is unknown, but several pathways are plausible. Parenting and affection display styles are heritable traits, and it may be that the genetic and environmental factors contributing to adverse parenting styles also contribute to autism trait emergence in toddlerhood (50, 51). Previous studies have suggested that parenting styles directly influence childhood behaviour, as children learn by repetition (52, 53). Parent-child relationships of children with childhood autism diagnoses are also more likely to be discordant than those of neurotypical offspring (54). This discordance is thought to be both a cause and consequence of difficulties in social understanding (55, 56), and it is possible that even at 18 months toddlers displaying more autism traits have greater difficulty relating to their parents, leading to greater discordance (57, 58). In support of this hypothesis a recent randomised controlled trial demonstrated that a 10-session therapist delivered parenting skills intervention, which promoted concordant interaction, led to a roughly 3 fold reduction in autism diagnoses 2 years later (59). However, parenting styles are at least partly heritable (60), hence it is also possible that the offspring of parents who naturally display more verbose and less collaborative parenting styles experience more difficulties developing social relationship abilities, and thus score more highly on the Q-CHAT. A final possibility is that maternal verbosity is in part a proxy measure of other forms of adversity: Verbosity has been previously shown to correlate with multiple measures of maternal stress (61), which in turn has previously been reported to correlate with a higher likelihood of offspring autism (62). All dimensions of parenting style are correlated with IMD rank in our data (Figure 1), and this is in keeping with a body of literature demonstrating associations between parenting style and socio-economic status (63). A more deeply phenotyped sample would be required to investigate how and if these different factors influence the relationship between maternal verbosity and Q-CHAT score. We do not seek to suggest that the emergence of autism traits is something parents can control, and a final possible interpretation of the correlation between maternal parenting style and autism trait emergence is reporting bias. Given that both the Q-CHAT and the parenting style questionnaire are self-reported tools individual patterns of response could relate to a wide number of factors, including mental state, intellectual ability and neurodevelopmental profile. Future studies could consider clinician administered measures to address this issue. There is another limitation to our findings here – we did not ask any questions about family composition or care arrangements beyond parent cohabiting status – we therefore do not know if the mother was the primary caregiver for each child included.

Based on previous literature, some of our results are expected, while others are unexpected. For instance, we showed that multiple measures of psychosocial disadvantage correlate with higher Q-CHAT scores. There is a significant body of evidence demonstrating that early life adversity affects several domains of early childhood behaviour, including cognitive (29), motor (30), and language (64) development, as well as emerging psychopathologies (25, 65). It is known that lower socio-economic status correlates with higher scores on the precursor to the Q-CHAT, the M-CHAT (32). Also, one previous study has specifically reported higher Q-CHAT scores in the offspring of depressed mothers (24). Therefore, our finding that maternal depressive symptom burden, measured using EPDS, correlates with offspring Q-CHAT score is not unexpected. In keeping with existing knowledge about neurodevelopment is our finding that two measures of social adversity correlate with higher Q-CHAT score. Our finding of a univariable association between maternal first language and Q-CHAT score is also in keeping with a body of previous literature which demonstrates a higher rate of autism diagnoses in children from immigrant backgrounds. It is likely that parent first language not being English represents an increased risk of experiencing other adversities (66), rather than inferring that being raised in a bilingual environment has an effect on autism trait emergence, which is not thought to be the case (67).

We unexpectedly found no association between sex and Q-CHAT score in any analysis performed. A handful of previous studies have demonstrated higher Q-CHAT scores in male toddlers compared to female toddlers, with small but significant average score differences (3.1 (68), 3.1 (69) and 1.9 (8)) reported. It is not immediately obvious why we do not see the same difference in our data, although it may be that in a larger sample this difference would have been apparent. Males in our cohort did in fact score 1.4 Q-CHAT points higher than females on average (Cohen’s d = 0.16), but the difference is not statistically significant. Similarly in a multivariable model the individual correlation between sex and Q-CHAT score is apparent (t=-2.32, p=0.020, Table 2) but did not survive FDR correction. It would be more appropriate to say that there is a trend towards males having higher Q-CHAT scores in our data than that there is no association at all.

We also found no significant association between parental history of ASD and Q-CHAT score in any analysis performed. A difference may reasonably have been expected based on the known familial increased likelihood of autism and ADHD diagnoses (48, 70). To date, one study has directly reported on the association between parental history of ASD and Q-CHAT score and found a large group difference, with the familial ASD history group having higher Q-CHAT scores at age 16-30 months (71). One other study has specifically examined the difference between Q-CHAT scores in individuals with and without an older sibling with autism, and also found significant group differences (72). It is not clear why we do not see the same effect here, although it is possible that the method in which we recorded family history (the mother was asked only if she or her partner had ever been diagnosed with autism) was too narrow a definition (a more broad dimensional assessment would have been preferrable), or alternatively it may be the case that we lacked sufficient positive cases (28 parents reported an ASD or ADHD diagnosis compared to 506 with no diagnosis) to have determinative power. Parents were also asked if the child participating in the study had an older sibling with an autism or ADHD diagnosis – as only 206 individuals had older siblings we have not included this variable in the main analysis. There was similarly no difference (t=-0.51, p=0.62) in mean Q-CHAT score between those with (n=23, mean Q-CHAT = 31.4) and without (n=183, mean Q-CHAT = 30.1) an older sibling with a neurodevelopmental diagnosis. This may again be due to an insufficient number of positive cases for determinant power.

It has been previously reported that preterm birth confers an increasing likelihood of both childhood autism diagnosis and greater early autism trait emergence (73, 74). One previous study reports Q-CHAT scores in a cohort of toddlers born before 30 weeks of gestation, who scored a mean of 33.7 (10), although to our knowledge no direct comparison of Q-CHAT scores in individuals born term and preterm has yet been presented. In our cohort we find no association between gestational age at birth and Q-CHAT score directly through univariable or multivariable associations, or indirectly via PCA latent components. One possibility is that early life autism trait emergence is less readily detected by screening tools in some preterm children (75, 76). Although we have used gestational age as a linear variable if we consider preterm birth as a binary variable there is also no difference between groups. The mean Q-CHAT of individuals born preterm is 30.1, and the mean Q-CHAT of individuals born at term is 30.1. The mean Q-CHAT scores in individuals born before 30 weeks gestation in our sample (n=36) is however 34.6, which is in keeping with the 33.7 average score reported by Wong et al. (2014) using the same criteria. Further research is needed to understand how the degree of prematurity effects early life autism trait emergence.

A finding of particular interest is how associations between demographic variables and Q-CHAT score were influenced by general neurodevelopment, which in our study is represented by BSID-III. All BSID-III composite scores correlated individually to the Q-CHAT score (Table 1). In a multivariable model without BSID-III scores four variables (two socio-demographic measures, and two measures of parenting style) were significantly associated with Q-CHAT score (Table 2). With BSID-III composite scores added to the model the two socio-demographic associations were no longer significant, although the BSID-III language composite score association was. This is possibly in part due to co-linearity of the input variables (Figure 1A). After transforming linear variables into latent orthogonal components with PCA, PC1 (associated with positive parenting styles and low socio-economic deprivation), was significantly negatively associated with Q-CHAT score with or without BSID-III scores as a confounder – i.e., more early life adversity was associated with more autism traits (Table 3). PC1 was significantly associated with Q-CHAT score in both models, suggesting that socio-demographic and parental factors are specifically influencing autism trait development as opposed to solely having a general effect on neurodevelopment. Using PC1, we can see how our original variables contribute to Q-CHAT score (Figure 1D). Some of the variables contributing to PC1 are expected based on our univariable results and previous literature; via PC1, early life adversity is associated with more autism traits, and maternal depression and more extreme parenting style is associated with more autism traits. Two variables however correlate in a less intuitive fashion. Firstly, maternal age is positively contributing to the association with Q-CHAT score (Figure 1D). This is not in keeping with a significant body of literature that suggests that the offspring of older parents have a higher likelihood of autism (77). One possible explanation is that there are aspects of social deprivation that we are not capturing with our variables, for example income or wider availability of family support, which may be related to both parental age and autism trait development. Secondly, maternal age at leaving full time education is positively contributing to the association with Q-CHAT score via the PC1. Given that greater social deprivation in our sample is in general associated with higher Q-CHAT scores this is somewhat counter-intuitive and is not in keeping with the one previous exploratory study to report on this association (24). There is a larger body of work regarding associations of parental education and childhood autism diagnoses, with at least some research suggesting that autism is more commonly diagnosed in the offspring of highly educated parents (78), so the same may be true of early life autism trait development. The reasons behind this difference are potentially complex, including greater access to medical professionals in more affluent families (79), diagnostic overshadowing (80) and more stigmatising views towards autism sometimes held by less educated parents (81).

Our findings suggest some possible avenues for future research. Deeply phenotyped and well powered prospective cohort studies of childhood autism are needed, but given the prevalence of the condition sample sizes would need to be extremely large to allow for firm conclusions to be drawn. A more logistically favourable approach to further examining some of the antecedents of autism trait development we (and other authors) have proposed would be to focus on groups hypothesised to be more likely to develop a high level of traits. This study design is well established when examining the sequelae of a family history of autism (82), and has also been used to study the effects of parental immigration (66) and depression (83). We suggest that a cohort experiencing severe psycho-social deprivation is a potential avenue in the study of early life autism traits.

## Limitations

There are some further limitations to our findings in addition to those discussed above. The cohort used is from a single study centre, and therefore may not be representative of the wider population. The sub-sample included in this study also differs from those excluded, in general experiencing less psycho-social adversity, with differences observed in IMD Rank, maternal parenting style and EPDS score. The nature of the scale is itself also a limitation: the Q-CHAT is parent rated, and therefore is indicative of the parent’s subjective assessment of their child, rather than an objective test (23); it is thus possible that reporting bias with common method variance could have altered our results.

A general linear model of all socio-demographic factors studied explained 20% of the variance of Q-CHAT score. Whilst this is a promising finding there are clearly a number of non-studied factors which may contribute to individual patterns of autism trait emergence, including genetics and medical comorbidities. Although emerging traits at age 18 months increase the likelihood of a future diagnosis of autism, the positive predictive value of a high Q-CHAT score (or indeed a high score on any early autism screening tool) is low (84). The prevalence of childhood autism in the UK is approximately 1.8% (85). If this prevalence is seen in our cohort then approximately 10 individuals may be expected to receive an autism diagnosis, meaning that what we are largely studying here are variations in the spectrum of typical development, which may (86) or may not (87) be of any real world relevance. Some of our more unexpected findings (for example the lack of a robust association between Q-CHAT score and sex) may in part be explained by a difference between the underlying nature of a clinical autism diagnoses and the expression of autism traits in the wider population. We hope in future to follow-up this cohort in childhood, which will allow us to re-analyse if the same factors we find here to be predictive of autism trait emergence are also predictive of diagnostic status.

## Conclusions

Autism traits at age 18 months in a typical population are associated with several prior exposures, most significantly parenting styles. In multivariable models 20% of variance of Q-CHAT score can be explained by socio-economic and parental factors, with the universal finding being that a less favourable environment results in a higher Q-CHAT score (more autism traits). Our results are of potential interest from two perspectives. Firstly, future authors investigating the Q-CHAT score and other measures of early autism traits should be aware of our findings as potential confounders or limiting factors in their work. Secondly, from our data it would be reasonable to expect a greater rate of diagnoses in more socio-economically deprived children, which does not currently occur. Are potential autism diagnoses being missed in more socially deprived groups?

## Declarations

### Ethical Approval

This project has received UK NHS research ethics committee approval (14/LO/1169, IRAS 138070), and conducted in accordance with the World Medical Association’s Code of Ethics (Declaration of Helsinki). Written informed consent was obtained from parents at recruitment into the study.

### Competing Interests

No author has a competing interest to declare.

### Data availability

The dHCP is an open-access project. Data from the project can be downloaded by registering at https://data.developingconnectome.org. Analyses presented here include data to be included in future releases.

### Author Contributions

All authors met ICJME criteria for authorship. **OGG** – conception, analysis, interpretation, writing original draft, final approval, accountability, **AC** – data collection, writing, review and editing, final approval, accountability, **SF** - data collection, writing, review and editing, final approval, accountability, **LF** – software, analysis, writing, review and editing, final approval, accountability, **SFM** – interpretation, writing, review and editing, final approval, accountability, **LH** – interpretation, writing, review and editing, final approval, accountability, **NH** – data curation, interpretation, writing, review and editing, final approval, accountability, **TC** – interpretation, writing, review and editing, final approval, accountability, **DM** – interpretation, analysis, writing, review and editing, final approval, accountability, **TA** – data collection, interpretation, writing, review and editing, final approval, accountability, **GM** – interpretation, analysis, writing, review and editing, final approval, accountability, **CN** – interpretation, analysis, writing, review and editing, final approval, accountability, **DE** – interpretation, analysis, final approval, accountability, **DB** – conception, analysis, interpretation, writing original draft, final approval, accountability.

### Funding

This work was supported by the European Research Council under the European Union’s Seventh Framework Programme (FP7/20072013)/ERC grant agreement no. 319456 (dHCP project). The authors acknowledge infrastructure support from the National Institute for Health Research (NIHR) Mental Health Biomedical Research Centre (BRC) at South London, Maudsley NHS Foundation Trust and King’s College London and the NIHR-BRC at Guys and St Thomas’ Hospitals NHS Foundation Trust (GSTFT). The authors also acknowledge support in part from the Wellcome Engineering and Physical Sciences Research Council (EPSRC) Centre for Medical Engineering at Kings College London [WT 203148/Z/16/Z], MRC strategic grant [MR/K006355/1], Medical Research Council Centre grant [MR/N026063/1], the Department of Health through an NIHR Comprehensive Biomedical Research Centre Award (to Guy’s and St. Thomas’ National Health Service (NHS) Foundation Trust in partnership with King’s College London and King’s College Hospital NHS Foundation Trust), the Sackler Institute for Translational Neurodevelopment at King’s College London and the European Autism Interventions (EU-AIMS) trial and the EU AIMS-2-TRIALS, a European Innovative Medicines Initiative Joint Undertaking under Grant Agreements No. 115300 and 777394, the resources of which are composed of financial contributions from the European Union’s Seventh Framework Programme (Grant FP7/2007–2013) and Horizon 2020 from the European Federation of Pharmaceutical industries and Associations companies’ in-kind contributions, and from Autism Speaks, Autistica, and the Simons Foundation for Autism Research Initiative. OGG is supported by a grant from the UK Medical Research Council / Sackler Foundation [MR/P502108/1]. SFM is supported by the UK Medical Research Council (MR/N013700/1) and King’s College London member of the MRC Doctoral Training Partnership in Biomedical Sciences. TA is supported by a MRC Clinician Scientist Fellowship [MR/P008712/1] and Transition Support Award [MR/V036874/1]. DB received support from a Wellcome Trust Seed Award in Science [217316/Z/19/Z]. The views expressed are those of the authors and not necessarily those of the NHS, the National Institute for Health Research or the Department of Health. The funders had no role in the design and conduct of the study; collection, management, analysis, and interpretation of the data; preparation, review, or approval of the manuscript; and decision to submit the manuscript for publication.

## Notes

### Competing Interest Statement

The authors have declared no competing interest.

### Summary of Updates

Figures altered, introduction and discussion substantially altered.

## References

1. Brett D, Warnell F, McConachie H, Parr JR. Factors Affecting Age at ASD Diagnosis in UK: No Evidence that Diagnosis Age has Decreased Between 2004 and 2014. J Autism Dev Disord. 2016;46(6):1974–84.

2. Daniels AM, Mandell DS. Explaining differences in age at autism spectrum disorder diagnosis: a critical review. Autism. 2014;18(5):583–97.

3. Crane L, Chester JW, Goddard L, Henry LA, Hill E. Experiences of autism diagnosis: A survey of over 1000 parents in the United Kingdom. Autism. 2016;20(2):153–62.

4. Schjølberg S, Shic F, Volkmar FR, Nordahl-Hansen A, Stenberg N, Torske T, et al. What are we optimizing for in autism screening? Examination of algorithmic changes in the M-CHAT. Autism Res. 2022;15(2):296–304.

5. Jullien S. Screening for autistic spectrum disorder in early childhood. BMC Pediatr. 2021;21(Suppl 1):349.

6. Toh T-H, Tan VW-Y, Lau PS-T, Kiyu A. Accuracy of Modified Checklist for Autism in Toddlers (M-CHAT) in detecting autism and other developmental disorders in community clinics. J Autism Dev Disord. 2018;48(1):28–35.

7. Guthrie W, Wallis K, Bennett A, Brooks E, Dudley J, Gerdes M, et al. Accuracy of Autism Screening in a Large Pediatric Network. Pediatrics. 2019;144(4).

8. Allison C, Baron-Cohen S, Wheelwright S, Charman T, Richler J, Pasco G, et al. The Q-CHAT (Quantitative CHecklist for Autism in Toddlers): a normally distributed quantitative measure of autistic traits at 18-24 months of age: preliminary report. J Autism Dev Disord. 2008;38(8):1414–25.

9. Ruta L, Chiarotti F, Arduino GM, Apicella F, Leonardi E, Maggio R, et al. Validation of the Quantitative Checklist for Autism in Toddlers in an Italian Clinical Sample of Young Children With Autism and Other Developmental Disorders. Front Psychiatry. 2019;10:488-.

10. Wong HS, Huertas-Ceballos A, Cowan FM, Modi N. Evaluation of early childhood social-communication difficulties in children born preterm using the Quantitative Checklist for Autism in Toddlers. J Pediatr. 2014;164(1):26–33.e1.

11. Magiati I, Goh DA, Lim SJ, Gan DZ, Leong JC, Allison C, et al. The psychometric properties of the Quantitative-Checklist for Autism in Toddlers (Q-CHAT) as a measure of autistic traits in a community sample of Singaporean infants and toddlers. Mol Autism. 2015;6:40.

12. Park S, Won EK, Lee JH, Yoon S, Park EJ, Kim AY. Reliability and Validity of the Korean Translation of Quantitative Checklist for Autism in Toddlers: A Preliminary Study. Soa Chongsonyon Chongsin Uihak. 2018;29(2):80–5.

13. Mohammadian M, Zarafshan H, Mohammadi MR, Karimi I. Evaluating Reliability and Predictive Validity of the Persian Translation of Quantitative Checklist for Autism in Toddlers (Q-CHAT). Iran J Psychiatry. 2015;10(1):64–70.

14. Allison C, Matthews FE, Ruta L, Pasco G, Soufer R, Brayne C, et al. Quantitative Checklist for Autism in Toddlers (Q-CHAT). A population screening study with follow-up: the case for multiple time-point screening for autism. BMJ Paediatrics Open. 2021;5(1):e000700.

15. Koolschijn PC, Geurts HM, van der Leij AR, Scholte HS. Are Autistic Traits in the General Population Related to Global and Regional Brain Differences? J Autism Dev Disord. 2015;45(9):2779–91.

16. Tsompanidis A, Aydin E, Padaigaitė E, Richards G, Allison C, Hackett G, et al. Maternal steroid levels and the autistic traits of the mother and infant. Mol Autism. 2021;12(1):51.

17. Thapar A, Rutter M. Genetic advances in autism. Journal of autism and developmental disorders. 2021;51:4321–32.

18. Chaste P, Leboyer M. Autism risk factors: genes, environment, and gene-environment interactions. Dialogues in clinical neuroscience. 2022.

19. Auyeung B, Ahluwalia J, Thomson L, Taylor K, Hackett G, O’Donnell KJ, et al. Prenatal versus postnatal sex steroid hormone effects on autistic traits in children at 18 to 24 months of age. Mol Autism. 2012;3(1):17.

20. Eldeeb SY, Ludwig NN, Wieckowski AT, Dieckhaus MF, Algur Y, Ryan V, et al. Sex differences in early autism screening using the Modified Checklist for Autism in Toddlers, Revised, with Follow-Up (M-CHAT-R/F). Autism. 2023:13623613231154728.

21. Gray PH, Edwards DM, O’Callaghan MJ, Gibbons K. Screening for autism spectrum disorder in very preterm infants during early childhood. Early Human Development. 2015;91(4):271–6.

22. Syn NL, Chan S-Y, Chia EWY, Ong WX, Phua D, Cai S, et al. Severity of nausea and vomiting in pregnancy and early childhood neurobehavioural outcomes: The Growing Up in Singapore Towards Healthy Outcomes study. Paediatric and Perinatal Epidemiology. 2021;35(1):98–108.

23. Ravi S, Chandrasekaran V, Kattimani S, Subramanian M. Maternal and birth risk factors for children screening positive for autism spectrum disorders on M-CHAT-R. Asian Journal of Psychiatry. 2016;22:17–21.

24. Goh DA, Gan D, Kung J, Baron-Cohen S, Allison C, Chen H, et al. Child, Maternal and Demographic Factors Influencing Caregiver-Reported Autistic Trait Symptomatology in Toddlers. J Autism Dev Disord. 2018;48(4):1325–37.

25. Kleine I, Vamvakas G, Lautarescu A, Falconer S, Chew A, Counsell SJ, et al. Postnatal maternal depressive symptoms and behavioural outcomes in term- and preterm-born toddlers. medRxiv. 2021:2021.09.21.21263881.

26. Schmengler H, El-Khoury Lesueur F, Yermachenko A, Taine M, Cohen D, Peyre H, et al. Maternal immigrant status and signs of neurodevelopmental problems in early childhood: The French representative ELFE birth cohort. Autism Res. 2019;12(12):1845–59.

27. Spencer-Smith MM, Spittle AJ, Lee KJ, Doyle LW, Anderson PJ. Bayley-III Cognitive and Language Scales in Preterm Children. Pediatrics. 2015;135(5):e1258–65.

28. Rao R, Trivedi S, Distler A, Liao S, Vesoulis Z, Smyser C, et al. Neurodevelopmental Outcomes in Neonates with Mild Hypoxic Ischemic Encephalopathy Treated with Therapeutic Hypothermia. Am J Perinatol. 2019;36(13):1337–43.

29. Ross GS, Perlman JM. Relationships of biological and environmental factors to cognition of preterm infants in the toddler and preschool periods. Dev Psychobiol. 2019;61(7):1100–6.

30. Ferreira L, Godinez I, Gabbard C, Vieira JLL, Caçola P. Motor development in school-age children is associated with the home environment including socio-economic status. Child Care Health Dev. 2018;44(6):801–6.

31. Neamah HH, Sudfeld C, McCoy DC, Fink G, Fawzi WW, Masanja H, et al. Intimate Partner Violence, Depression, and Child Growth and Development. Pediatrics. 2018;142(1).

32. Khowaja MK, Hazzard AP, Robins DL. Sociodemographic Barriers to Early Detection of Autism: Screening and Evaluation Using the M-CHAT, M-CHAT-R, and Follow-Up. J Autism Dev Disord. 2015;45(6):1797–808.

33. Harker CM, Ibañez LV, Nguyen TP, Messinger DS, Stone WL. The Effect of Parenting Style on Social Smiling in Infants at High and Low Risk for ASD. J Autism Dev Disord. 2016;46(7):2399–407.

34. Carter AS, Martínez-Pedraza Fde L, Gray SA. Stability and individual change in depressive symptoms among mothers raising young children with ASD: maternal and child correlates. J Clin Psychol. 2009;65(12):1270–80.

35. Rayport YK, Sania A, Lucchini M, Du Plessis C, Potter M, Springer PE, et al. Associations of adverse maternal experiences and diabetes on postnatal maternal depression and child social-emotional outcomes in a South African community cohort. PLOS Glob Public Health. 2022;2(10):e0001124.

36. Hines M, Carpenito T, Martens A, Iizuka A, Aspinwall B, Zimmerman E. The home environment and its relation to vocalizations in the first year of life. Pediatr Med. 2022;5.

37. Bayley N. Bayley scales of infant and toddler development, third edition. San Antonio, TX: Harcourt; 2006.

38. Wolke D, Jaekel J, Hall J, Baumann N. Effects of sensitive parenting on the academic resilience of very preterm and very low birth weight adolescents. J Adolesc Health. 2013;53(5):642–7.

39. Vanes LD, Hadaya L, Kanel D, Falconer S, Ball G, Batalle D, et al. Associations Between Neonatal Brain Structure, the Home Environment, and Childhood Outcomes Following Very Preterm Birth. Biological Psychiatry Global Open Science. 2021;1(2):146–55.

40. Arnold DS, O’leary SG, Wolff LS, Acker MM. The Parenting Scale: a measure of dysfunctional parenting in discipline situations. Psychological assessment. 1993;5(2):137.

41. Cox JL, Holden JM, Sagovsky R. Detection of postnatal depression: development of the 10-item Edinburgh Postnatal Depression Scale. The British journal of psychiatry. 1987;150(6):782–6.

42. Benjamini Y, Hochberg Y. Controlling the False Discovery Rate: A Practical and Powerful Approach to Multiple Testing. Journal of the Royal Statistical Society: Series B (Methodological). 1995;57(1):289–300.

43. Jolliffe IT, Cadima J. Principal component analysis: a review and recent developments. Philosophical Transactions of the Royal Society A: Mathematical, Physical and Engineering Sciences. 2016;374(2065):20150202.

44. Murray M, Bloom, J. FDRestimation: Estimate, Plot, and Summarize False Discovery Rates,. R Package. 1.0.1 ed2020.

45. Wei T, Simko, V. R package ‘corrplot’: Visualization of a Correlation Matrix. 0.92 ed2021.

46. Sun Z, Yang L, Bai X, Du W, Shen G, Fei J, et al. Maternal ambient air pollution exposure with spatial-temporal variations and preterm birth risk assessment during 2013–2017 in Zhejiang Province, China. Environment International. 2019;133:105242.

47. Werling DM, Geschwind DH. Sex differences in autism spectrum disorders. Curr Opin Neurol. 2013;26(2):146–53.

48. Miller M, Musser ED, Young GS, Olson B, Steiner RD, Nigg JT. Sibling Recurrence Risk and Cross-aggregation of Attention-Deficit/Hyperactivity Disorder and Autism Spectrum Disorder. JAMA Pediatr. 2019;173(2):147–52.

49. Agrawal S, Rao SC, Bulsara MK, Patole SK. Prevalence of Autism Spectrum Disorder in Preterm Infants: A Meta-analysis. Pediatrics. 2018;142(3):e20180134.

50. Shaw ZA, Starr LR. Intergenerational transmission of emotion dysregulation: The role of authoritarian parenting style and family chronic stress. Journal of Child and Family Studies. 2019;28:3508–18.

51. Klahr AM, Burt SA. Elucidating the etiology of individual differences in parenting: A meta-analysis of behavioral genetic research. Psychol Bull. 2014;140(2):544–86.

52. Johnston C, Murray C, Hinshaw SP, Pelham WE, Hoza B. Responsiveness in interactions of mothers and sons with ADHD: Relations to maternal and child characteristics. Journal of abnormal child psychology. 2002;30(1):77–88.

53. Miller-Lewis LR, Baghurst PA, Sawyer MG, Prior MR, Clark JJ, Arney FM, et al. Early childhood externalising behaviour problems: Child, parenting, and family-related predictors over time. Journal of abnormal child psychology. 2006;34(6):886–901.

54. Crowell JA, Keluskar J, Gorecki A. Parenting behavior and the development of children with autism spectrum disorder. Comprehensive psychiatry. 2019;90:21–9.

55. Ventola P, Lei J, Paisley C, Lebowitz E, Silverman W. Parenting a Child with ASD: Comparison of Parenting Style Between ASD, Anxiety, and Typical Development. J Autism Dev Disord. 2017;47(9):2873–84.

56. Gau SS, Chou MC, Lee JC, Wong CC, Chou WJ, Chen MF, et al. Behavioral problems and parenting style among Taiwanese children with autism and their siblings. Psychiatry Clin Neurosci. 2010;64(1):70–8.

57. Craig F, Operto FF, De Giacomo A, Margari L, Frolli A, Conson M, et al. Parenting stress among parents of children with Neurodevelopmental Disorders. Psychiatry Res. 2016;242:121–9.

58. Wan MW, Green J, Scott J. A systematic review of parent-infant interaction in infants at risk of autism. Autism. 2019;23(4):811–20.

59. Whitehouse AJO, Varcin KJ, Pillar S, Billingham W, Alvares GA, Barbaro J, et al. Effect of Preemptive Intervention on Developmental Outcomes Among Infants Showing Early Signs of Autism: A Randomized Clinical Trial of Outcomes to Diagnosis. JAMA Pediatrics. 2021;175(11):e213298-e.

60. Oliver BR, Trzaskowski M, Plomin R. Genetics of parenting: The power of the dark side. Dev Psychol. 2014;50(4):1233–40.

61. McQuillan ME, Bates JE, Staples AD, Deater-Deckard K. Maternal stress, sleep, and parenting. Journal of Family Psychology. 2019;33(3):349.

62. Khambadkone SG, Cordner ZA, Tamashiro KLK. Maternal stressors and the developmental origins of neuropsychiatric risk. Front Neuroendocrinol. 2020;57:100834.

63. La Placa V, Corlyon J. Unpacking the relationship between parenting and poverty: Theory, evidence and policy. Social Policy and Society. 2016;15(1):11–28.

64. Wild KT, Betancourt LM, Brodsky NL, Hurt H. The effect of socio-economic status on the language outcome of preterm infants at toddler age. Early Hum Dev. 2013;89(9):743–6.

65. de Laat SAA, Huizink AC, Hof MH, Vrijkotte TGM. Socio-economic inequalities in psychosocial problems of children: mediating role of maternal depressive symptoms. Eur J Public Health. 2018;28(6):1062–8.

66. Abdullahi I, Wong K, Bebbington K, Mutch R, de Klerk N, Cherian S, et al. Diagnosis of Autism Spectrum Disorder According to Maternal-Race Ethnicity and Country of Birth: A Register-Based Study. J Autism Dev Disord. 2019;49(9):3611–24.

67. Kašćelan D, Katsos N, Gibson JL. Relations Between Bilingualism and Autistic-Like Traits in a General Population Sample of Primary School Children. Journal of Autism and Developmental Disorders. 2019;49(6):2509–23.

68. Kung KT, Constantinescu M, Browne WV, Noorderhaven RM, Hines M. No relationship between early postnatal testosterone concentrations and autistic traits in 18 to 30-month-old children. Mol Autism. 2016;7:15.

69. Auyeung B, Taylor K, Hackett G, Baron-Cohen S. Foetal testosterone and autistic traits in 18 to 24-month-old children. Mol Autism. 2010;1(1):11.

70. Sandin S, Lichtenstein P, Kuja-Halkola R, Larsson H, Hultman CM, Reichenberg A. The Familial Risk of Autism. JAMA. 2014;311(17):1770–7.

71. Ben-Sasson A, Robins DL, Yom-Tov E. Risk Assessment for Parents Who Suspect Their Child Has Autism Spectrum Disorder: Machine Learning Approach. J Med Internet Res. 2018;20(4):e134-e.

72. Pasco G, Davies K, Ribeiro H, Tucker L, Allison C, Baron-Cohen S, et al. Comparison of Parent Questionnaires, Examiner-Led Assessment and Parents’ Concerns at 14 Months of Age as Indicators of Later Diagnosis of Autism. J Autism Dev Disord. 2021;51(3):804-13.

73. Crump C, Sundquist J, Sundquist K. Preterm or Early Term Birth and Risk of Autism. Pediatrics. 2021;148(3).

74. Guy A, Seaton SE, Boyle EM, Draper ES, Field DJ, Manktelow BN, et al. Infants born late/moderately preterm are at increased risk for a positive autism screen at 2 years of age. The Journal of pediatrics. 2015;166(2):269–75. e3.

75. Gray PH. M-CHAT autism screening may be inaccurate among toddlers born very preterm. The Journal of Pediatrics. 2017;182:401–4.

76. Moore T, Johnson S, Hennessy E, Marlow N. Screening for autism in extremely preterm infants: problems in interpretation. Developmental Medicine & Child Neurology. 2012;54(6):514–20.

77. Parner ET, Baron-Cohen S, Lauritsen MB, Jørgensen M, Schieve LA, Yeargin-Allsopp M, et al. Parental age and autism spectrum disorders. Ann Epidemiol. 2012;22(3):143–50.

78. King MD, Bearman PS. Socio-economic status and the increased prevalence of autism in California. American sociological review. 2011;76(2):320–46.

79. Winter AS, Fountain C, Cheslack-Postava K, Bearman PS. The social patterning of autism diagnoses reversed in California between 1992 and 2018. Proceedings of the National Academy of Sciences. 2020;117(48):30295–302.

80. Avlund SH, Thomsen PH, Schendel D, Jørgensen M, Carlsen AH, Clausen L. Factors Associated with a Delayed Autism Spectrum Disorder Diagnosis in Children Previously Assessed on Suspicion of Autism. Journal of Autism and Developmental Disorders. 2021;51(11):3843–56.

81. Azim A, Rdesinski RE, Phelps R, Zuckerman KE. Nonclinical factors in autism diagnosis: Results from a national health care provider survey. Journal of Developmental & Behavioral Pediatrics. 2020;41(6):428–35.

82. Bolton P, Macdonald H, Pickles A, Rios Pa, Goode S, Crowson M, et al. A case-control family history study of autism. Journal of child Psychology and Psychiatry. 1994;35(5):877–900.

83. Chen L-C, Chen M-H, Hsu J-W, Huang K-L, Bai Y-M, Chen T-J, et al. Association of parental depression with offspring attention deficit hyperactivity disorder and autism spectrum disorder: A nationwide birth cohort study. Journal of affective disorders. 2020;277:109–14.

84. Thabtah F, Peebles D. Early autism screening: a comprehensive review. International journal of environmental research and public health. 2019;16(18):3502.

85. Roman-Urrestarazu A, van Kessel R, Allison C, Matthews FE, Brayne C, Baron-Cohen S. Association of Race/Ethnicity and Social Disadvantage With Autism Prevalence in 7 Million School Children in England. JAMA Pediatrics. 2021;175(6):e210054-e.

86. Mottron L. A radical change in our autism research strategy is needed: Back to prototypes. Autism Research. 2021;n/a(n/a).

87. Constantino JN. Response to “A Radical Change in Our Autism Research Strategy is Needed: Back to Prototypes” by Mottron et al. (2021). Autism Research. 2021;n/a(n/a).

