## Supplementary for "Clinical, socio-demographic, and parental correlates of early autism traits in a community cohort"

### Supplementary Material

***Supplementary Table S1 –*** *Comparison of individuals included (n=536) and excluded (n=108) from analysis. Group differences assessed by t-test or chi-squared test as appropriate.*

| | **Mean Included** | **Mean Excluded** | **t** | **p** |
|---|---|---|---|---|
| Total Q-CHAT Score | 30.1 | 31.2 | 1.00 | 0.318 |
| Gestational Age at Birth [Weeks] | 38.2 | 38.9 | 1.74 | 0.083 |
| BMI | 24.7 | 24.1 | 0.96 | 0.334 |
| Mother Age [Years] | 33.9 | 34.3 | -0.88 | 0.377 |
| | **M/F Included** | **M/F Excluded** | **$\chi^2$** | **p** |
| Sex | 278/258 | 65/43 | 2.17 | 0.140 |
| | **Y/N Included** | **Y/N Excluded** | | |
| Parent ASD/ADHD Diagnosis | 28/508 | 5/96 | 0.13 | 0.990 |
| | **Mean Included** | **Mean Excluded** | **t** | **p** |
| IMD Rank | 14588 | 11712 | **-3.96** | **<0.001** |
| CSPS | 20.5 | 20.5 | -0.08 | 0.932 |
| Mother Education | 23.2 | 23.6 | -0.86 | 0.388 |
| | **Y/N Included** | **Y/N Excluded** | **$\chi^2$** | **P** |
| Mother 1st Language English | 338/198 | 55/49 | 3.38 | 0.065 |
| | **Y/N Included** | **Y/N Excluded** | | |
| Parents Cohabiting | 520/16 | 103/15 | 0.337 | 0.561 |
| | **Mean Included** | **Mean Excluded** | **t** | **P** |
| Mother Laxness | 2.9 | 3.2 | **2.71** | **0.007** |
| Mother Overreactivity | 2.2 | 2.3 | 1.46 | 0.142 |
| Mother Verbosity | 3.4 | 3.6 | **2.25** | **0.025** |
| Mother EPDS | 4.5 | 2.6 | **-3.31** | **0.001** |

***Supplementary Table S2*** *– Correlates between PC1-3 and linear variables. PC1 is associated with positive parenting styles and low socioeconomic deprivation, PC2 is associated with low socioeconomic deprivation with expressive parenting styles, and PC3 is associated with clinical adversity.*

| | **PC1** | | **PC2** | | **PC3** | |
|---|---|---|---|---|---|---|
| | **r** | **p** | **r** | **p** | **r** | **p** |
| Gestational Age at Birth | -0.07 | 0.083 | 0.09 | 0.304 | **-0.69** | **<0.001*** |
| BMI | -0.01 | 0.886 | **-0.13** | **0.017*** | **0.49** | **<0.001*** |
| Mother Age | **0.25** | **<0.001*** | **0.72** | **<0.001*** | **0.19** | **<0.001*** |
| IMD Rank | **0.31** | **<0.001*** | **0.45** | **<0.001*** | -0.01 | 0.783 |
| CSPS | **0.39** | **<0.001*** | **0.30** | **<0.001*** | **0.40** | **<0.001*** |
| Mother Education | -0.08 | 0.427 | **0.63** | **<0.001*** | **-0.28** | **<0.001*** |
| Mother Laxness | **-0.69** | **<0.001*** | **0.10** | **0.019*** | 0.07 | 0.084 |
| Mother Overreactivity | **-0.57** | **<0.001*** | **0.27** | **<0.001*** | **0.14** | **0.001*** |
| Mother Verbosity | **-0.73** | **<0.001*** | **0.20** | **<0.001*** | **0.11** | **0.008*** |
| Mother EPDS | **-0.29** | **<0.001*** | 0.01 | 0.819 | **0.20** | **<0.001*** |

**p value remains significant after FDR multiple comparison correction*

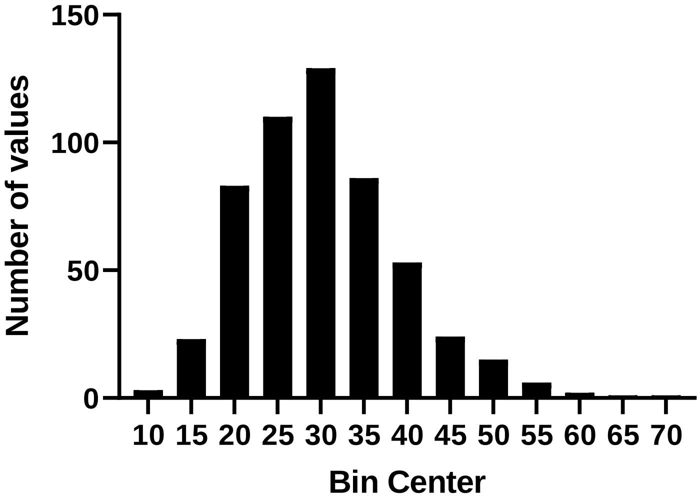


***Supplementary Figure S1 –*** *Frequency Distribution of Q-CHAT scores. Shapiro Wilk W=0.973, p<0.001. 90/536 individuals scored 39 (the cut-off proposed for further investigation by Allison et al, 2021) or greater.*
